# Polycomb Group Ring Finger Protein 6 suppresses Myc-induced lymphomagenesis

**DOI:** 10.1101/2021.12.02.470967

**Authors:** Nina Tanaskovic, Mattia Dalsass, Giorgia Ceccotti, Marco Filipuzzi, Alessandro Verrecchia, Paola Nicoli, Mirko Doni, Daniela Olivero, Diego Pasini, Haruhiko Koseki, Arianna Sabò, Andrea Bisso, Bruno Amati

## Abstract

Max is an obligate dimerization partner for the Myc transcription factors and for several repressors, such as Mnt, Mxd1-4 and Mga, collectively thought to antagonize Myc function in transcription and oncogenesis. Mga, in particular, is part of the variant Polycomb group repressive complex PRC1.6. Here, we show that ablation of the distinct PRC1.6 subunit Pcgf6 – but not Mga – accelerates Myc-induced lymphomagenesis in Eµ-*myc* transgenic mice. Unexpectedly, however, Pcgf6 loss shows no significant impact on transcriptional profiles, in neither pre-tumoral B-cells, nor lymphomas. Altogether, these data unravel an unforeseen, Mga- and PRC1.6-independent tumor suppressor activity of Pcgf6.

## Introduction

The Myc-Max network is constituted by a set of transcription factors that dimerize and bind DNA via a common basic-helix-loop-helix-leucine zipper motif (bHLH-LZ). Max is a key node in this network, acting as an obligate dimerization partner for proteins of the Myc (c-, N- and L-Myc) and Mxd/Mga subfamilies (Mxd1-4, Mnt and Mga), which activate and repress transcription, respectively, by binding to the same consensus DNA element, the E-box CACGTG and variants thereof (Carroll et al. 2018). Mxd1-4 and Mnt share a short N-terminal domain responsible for recruitment of mSin3/HDAC corepressor complexes. Mga lacks this domain, but was independently identified - together with Max - as a component of the variant Polycomb group (PcG) repressive complex PRC1.6, characterized by the presence of two distinct PcG- and E2F-family proteins (respectively, Pcgf6 and E2f6) (Ogawa et al. 2002; Gao et al. 2012; Carroll et al. 2018; Llabata et al. 2020). In mouse embryonic stem cells (mESCs), depletion of Mga led to dissociation of other PRC1.6 subunits (Pcgf6, E2f6 and L3mbtl2) from chromatin (Endoh et al. 2017; Stielow et al. 2018; Scelfo et al. 2019). Altogether, these findings suggest that Mga/Max and the associated PRC1.6 complex may counteract transcriptional activation by Myc and E2F at common target genes, and may thus also antagonize their growth-promoting and oncogenic activities.

A number of observations pointed to a tumor suppressor function of the Mga/Max dimer. First, genome sequencing studies revealed loss-of-function mutations in Mga in a wide variety of tumors (Schaub et al. 2018). Loss of Max was also observed, but appears to be restricted to neuroendocrine tumors, including pheochromocytoma (PC) (Comino-Mendez et al. 2011; Burnichon et al. 2012) and small-cell lung cancer (SCLC) (Romero et al. 2014; Llabata et al. 2021). In SCLC, mutations affecting the different network members (Max loss; Mga loss; Myc amplification) occur in a mutually exclusive manner, pointing to a common functional consequence (Romero et al. 2014). Formal evidence for this hypothesis was provided in two SCLC mouse models, in which deletion of Max could either abrogate tumorigenesis if combined with a *MYCL* transgene, or favor it following loss of the Rb1 and Trp53 tumors suppressors (Augert et al. 2020). Hence, in neuroendocrine tumors loss of Mga/Max/PRC1.6 repressor function may be sufficient to bypass the requirement for Myc activity, as recently shown in Max-null human SCLC cell lines (Llabata et al. 2021). In other lineages, the essential role of Max as a Myc partner (Amati et al. 1993) may prevent its loss, but may still co-exist with its antagonist activities in complex with either Mga or Mxd/Mnt proteins. In line with these observations, loss of Mga in a murine Myc-proficient non-small cell lung cancer model accelerated tumor growth and caused de-repression of PRC1.6, E2F and Myc/Max target genes (Mathsyaraja et al. 2021).

One of the tumor types with recurrent mutations in Mga is diffuse large B-cell lymphoma (DLBCL) (Reddy et al. 2017; Lee et al. 2020), in which Mga also scored as a top hit in a reverse-genetic screen for tumor suppressor genes (Reddy et al. 2017). In Eµ-*myc* transgenic mice, a widely used model for Myc-driven B-cell lymphoma, Max was essential for lymphomagenesis (Mathsyaraja et al. 2019); more surprisingly, Mnt also showed tumor-promoting activity in this model, owing most likely to selective suppression of Myc-induced apoptosis (Campbell et al. 2017; Nguyen et al. 2020). Here, we addressed whether loss of either Mga or the PRC1.6-restricted subunit Pcgf6 (Gao et al. 2012) potentiate lymphomagenesis in Eµ-*myc* mice. Unexpectedly, our data point to a distinct function of Pcgf6 in tumor suppression, independent from either Mga, PRC1.6 or transcriptional control.

## Results & Discussion

### Loss of Pcgf6 accelerates Myc-induced lymphomagenesis

To address the roles of Mga and Pcgf6 in Myc-induced lymphomagenesis, we combined the Eμ-*myc* (Adams et al. 1985) and CD19-*Cre* transgenes (Rickert et al. 1997) – thus expressing both Myc and Cre recombinase from the pro B-cell stage – with either the conditional knockout alleles *Mga*^*fl*^ (Washkowitz et al. 2015) or *Pcgf6*^*fl*^ (Endoh et al. 2017) (Supplemental Table S1). While targeting *Mga* showed no effect (Fig. S1A-C), deletion of *Pcgf6* significantly enhanced Eμ-*myc*-dependent lymphomagenesis, with *Pcgf6*^*+/fl*^ and *Pcgf6*^*fl/fl*^ animals showing progressive reductions in median disease-free survival, and increased disease penetrance (Fig. 1A). Relative to *Pcgf6*^*+/+*^ controls, *Pcgf6*^*+/fl*^ and *Pcgf6*^*fl/fl*^ tumors (hereafter *Pcgf6*^*+/*Δ^ and *Pcgf6*^Δ/Δ^ or KO) showed proportionate decreases in *Pcgf6* mRNA levels (Fig. 1B), and immunoblot analysis confirmed loss of the protein in the latter (Fig. 1C). The *Pcgf6* genotype affected neither the differentiation stage of the tumors, with comparable proportions arising from naive mature B-cells (B220^+^ IgM^+^) and B-cell precursors (B220^+^ IgM^-^) (Fig. 1D) (Langdon et al. 1986), nor their pathological classification, all examined cases showing DLBCL/Burkitt’s like features (Supplemental Table S2). Finally, we exploited our RNA-seq profiles (see below) to analyze tumor clonality (Barbosa et al. 2020): in contrast with the concept that lymphomas arising in Eµ-*myc* mice are monoclonal, as classically determined by PCR (Adams et al, 1985), we detected multiple clones in most tumors, regardless of their *Pcgf6* genotype (Supplemental Fig. S2; Table S3); most relevant here, accelerated tumor onset in the *Pcgf6*^*+/f*^ and *Pcgf6*^*f/f*^ backgrounds could not simply be ascribed to increased clonality. Altogether, we conclude that Pcgf6, functions as a dose-dependent, haplo-insufficient tumor suppressor in Myc-induced lymphomagenesis, without altering the gross pathological and cellular features of the resulting tumors. Unlike *Pcgf6, Mga* showed no tumor suppressor activity in Eµ-*myc* mice, pointing to a PRC1.6-independent function of Pcgf6 in this model.

**Figure 1.**
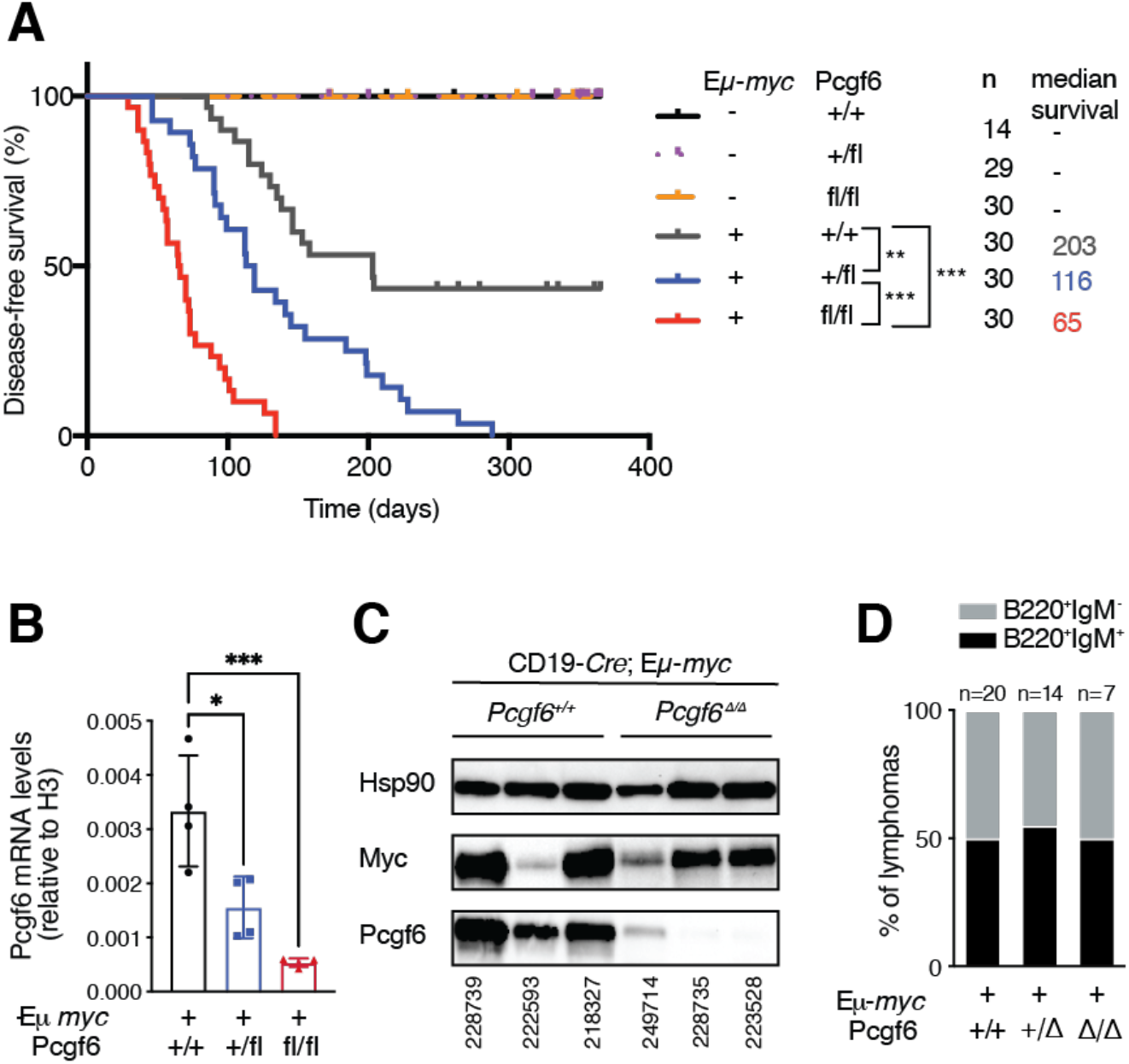
Loss of *Pcgf6* cooperates with Myc overexpression in B-cell lymphoma development. **A**. Disease-free survival curves for mice of the indicated Eµ-*myc* and *Pcgf6* genotypes (all with the CD19-*Cre* transgene). The number of mice (n) and median survival (in days) are indicated. **B**. RT-qPCR performed on the mRNA extracted from sorted CD19^+^ B-cells from infiltrated lymph nodes of CD19-*Cre*; Eμ-*myc*; *Pcgf6*^*fl/fl*^ mice. * p<0.05; ** p<0.001; *** p < 0.0001; **C**. Western blot analysis of Pcgf6 and Myc protein expression in infiltrated lymph nodes from either CD19-*Cre*; Eµ-*myc*; *Pcgf6*^*+/+*^ or CD19-*Cre*; Eµ-*myc*; *Pcgf6*^Δ/Δ^ tumors. Hsp90 was used as loading control. One representative mouse per genotype is shown and mice IDs are indicated at the bottom. Note that a residual Pcgf6 signal in *Pcgf6*^Δ/Δ^ samples might be due to infiltrating non-deleted cells. **D**. Immunophenotyping of B220 and IgM reveals similar proportions of B220^+^ IgM^+^ and B220^+^ IgM^-^ tumors among Eµ-*myc* lymphomas of the indicated *Pcgf6* genotypes. The numbers above each bar represent number of mice analyzed for each genotype.

### Loss of Pcgf6 affects Myc-induced apoptosis in B-cells

Young Eµ-*myc* mice show a characteristic expansion of pre-tumoral B-cells, counter-balanced by a concomitant increase in apoptosis (Nilsson et al. 2005). Monitoring of bone marrow B220^+^CD19^+^ B-cells revealed that their fraction was significantly increased in the *Pcgf6*^*f/f*^ background (Fig. 2A) correlating with an impairment in Myc-induced apoptosis (Fig. 2B). In contrast with the effect on apoptosis, loss of Pcgf6 caused no major alterations in the cell cycle profiles of B220^+^CD19^+^ B-cells, in either control or Eµ-*myc* transgenic mice (Fig. 2C). Altogether, our data suggest that the accelerated lymphoma onset in Eµ-*myc*; CD19-*Cre*; *Pcgf6*^*fl/fl*^ mice may be explained – at least in part – by increased survival at the pre-tumoral stage, which might favor the expansion of the B220^+^CD19^+^ B cell pool, thus increasing the opportunities for the emergence of tumor clones.

**Figure 2.**
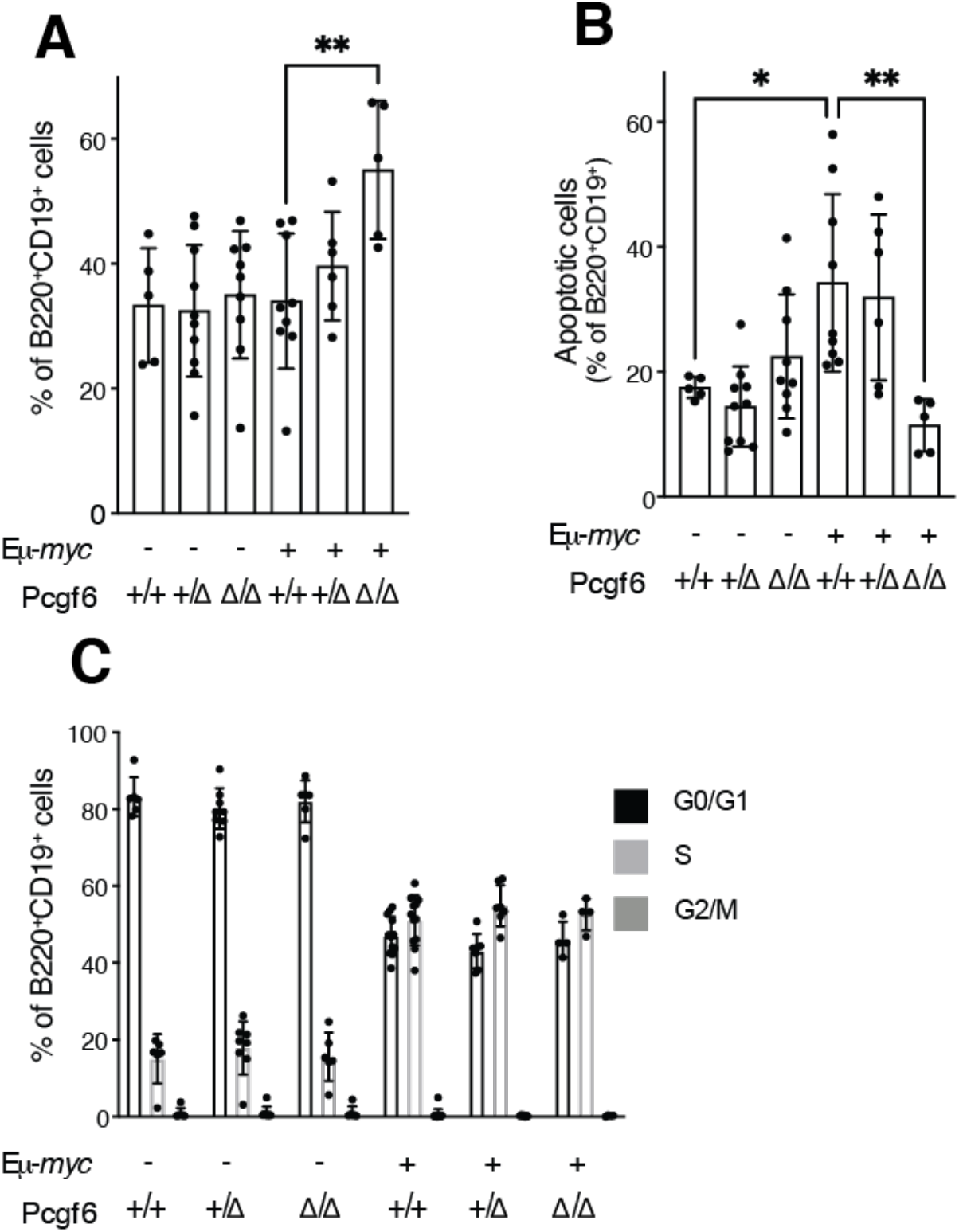
Pcgf6 loss affects Myc-induced apoptosis, but not proliferation in bone marrow B-cells. **A**. Fraction of B220^+^ CD19^+^ cells in the bone marrow (BM) of the indicated experimental groups. **B**. Fraction of apoptotic B220^+^ CD19^+^ cells, based on Red-VAD-FMK staining for caspase activity. **C**. Fraction of B220^+^ CD19^+^ cells in the G0/G1, S and G2/M phases of the cell cycle, as determined by EdU and Hoechst staining. * p<0.05; ** p<0.001

### Loss of Pcgf6 does not affect Myc-dependent transcription

As assayed by RNA-seq profiling, pre-tumoral Eμ-*myc* B-cells (P) show characteristic changes in gene expression relative to control non-transgenic B-cells (C) (Sabò et al. 2014). This was confirmed in our cohorts, with separate clustering of the C and P samples; within each cluster, however, the *Pcgf6* genotypes remained intermingled (Fig. 3A). At either the C or P stage, calling for differentially expressed genes (DEGs) yielded virtually no changes between the WT and KO samples (Supplemental Table S4). Taking WT B-cells as a common control, similar numbers of DEGs were called in the pre-tumoral samples (P), with a large overlap between the *Pcgf6* WT and KO genotypes (Fig. 3B-D, Supplemental Table S4). Similarly, RNA-seq profiling of tumor samples (T) yielded high correlation indices among all tumors, no clustering according to *Pcgf6* status, similar transcriptional changes in the KO and WT tumors relative to control B-cells, and very few DEGs (78 up and 65 down) in KO relative to WT tumors (Supplemental Fig. S3A-D). Most noteworthy here, while *Pcgf6* was not called as DEG in this comparison, the RNA-seq profiles confirmed the complete absence of *Pcgf6* exons 2 and 3 in KO tumors (Supplemental Fig. S3E). In conclusion, Pcgf6 impacted neither on steady-state gene expression profiles, nor on Myc-dependent responses during B-cell lymphomagenesis.

**Figure 3.**
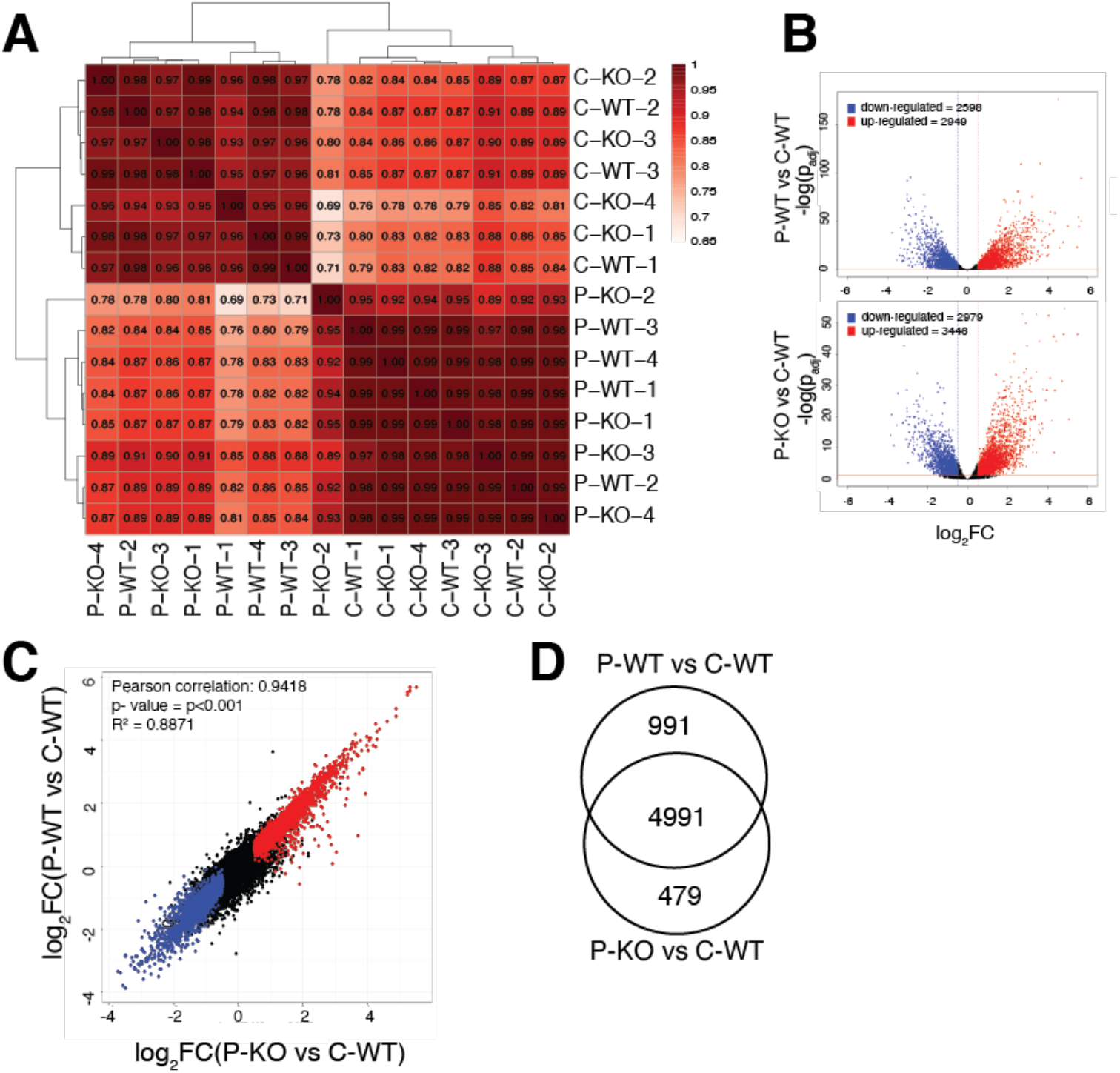
Pcgf6 loss does not affect Myc-dependent transcription. RNA-seq profiles were generated from control non-transgenic and pre-tumoral Eµ-*myc* B-cells (labeled C and P, respectively) with either *Pcgf6*^*+/*Δ^ (WT) or *Pcgf6*^Δ/Δ^ (KO) genotypes. All samples are indicated by the stage (C or P) followed by the Pcgf6 genotype (WT or KO) and the sample number. For C-WT, n=3; C-KO, P-WT and P-KO, n=4 **A**. Pairwise Pearson correlation between all samples, based on their RNA-seq profiles. **B**. Volcano plots showing the differentially expressed genes (DEGs) called in P-WT (top) or P-KO (bottom) with C-WT as a common control. The horizontal and vertical lines within the plots mark the statistical criteria used for calling DEGs, indicating the thresholds for significance (p_adj_<0.05) and fold change (FC) (|log_2_FC>0.5|). Up- and down-regulated DEGs are shown in red and blue, respectively, and their numbers indicated at the top. All DEGs are listed in Supplemental Table S4. **C**. Comparison of the DEGs called in P-WT (Y-axis) and P-KO (X-axis), as defined in B. The DEGs are colored based on the call in the x-axis. R^2^=0.8871 represents the coefficient of determination. **D**. Venn diagram showing the overlap in DEGs called in P-WT and P-KO.

### Mga-dependent recruitment of Pcgf6 to active chromatin

At first sight, the limited impact of Pcgf6 on transcriptional profiles appears at odds with its known function as a component of the PRC1.6 complex. The latter should depend on Mga, which is required both for the integrity of PRC1.6 and for the recruitment of Pcgf6 to chromatin, as shown in mouse embryonic stem cells (mESCs) and lung tumor cells (Gao et al. 2012; Endoh et al. 2017; Stielow et al. 2018; Scelfo et al. 2019; Mathsyaraja et al. 2021). To address the status of PRC1.6 in our lymphomas, we derived primary lymphoma cultures from Eµ-*myc* control mice and their *Mga*^*-/-*^ and *Pcgf6*^*-/-*^ counterparts (Supplemental Fig. S4A). We then used these cells for ChIP-seq profiling of Pcgf6, alongside active histone marks (H3K4me3, H3K4me1, H3K27ac), as well as the PRC2- and PRC1-associated repressive marks H3K27me3 and H2AK119Ub (Di Croce and Helin 2013). Analysis of the distribution of ChIP-seq reads among annotated promoters and distal sites in the genome (Supplemental Fig. S4B), revealed several key features. First, the Pcgf6 signal observed in the control Eµ-*myc* lymphoma was lost not only in *Pcgf6*^*-/-*^, but also in *Mga*^*-/-*^ cells, in line with the role of Mga in recruiting Pcgf6 to chromatin. Second, Pcfg6 did not co-localize with the PRC-associated marks H3K27me3 and H2AK119Ub, but showed preferential binding to active chromatin, as previously observed in mESCs (Stielow et al. 2018; Scelfo et al. 2019).

It is noteworthy here that the propensity to widely associate with active regulatory elements (promoters and enhancers) is a fundamental feature shared between Myc/Max and Mga/Max/PRC1.6 complexes (Gao et al. 2012; Sabò et al. 2014; Kress et al. 2015; Endoh et al. 2017; Stielow et al. 2018; Scelfo et al. 2019; Mathsyaraja et al. 2021). As recently documented for Myc, this initial non-specific engagement on DNA is a prerequisite for sequence (i.e. E-box) recognition and selective gene regulation (Pellanda et al. 2021). Owing to the small number of Pcgf6*-* and Mga-null lymphoma cell lines available in our work, and to the limiting availability of compound Eµ-*myc; Pcfg6*^Δ/Δ^ mice (Supplemental Table S1) precluding ChIP-seq analysis of pre-tumoral B-cells (Sabò et al. 2014), we were unable to address whether loss of PRC1.6 activity may significantly impact Myc’s binding profiles in our model. This scenario appears unlikely, however, given that Pcgf6 loss showed no significant impact on Myc-associated gene expression profiles. Altogether, while Pcgf6 shows Mga-dependent DNA binding, as expected in the context of the PRC1.6 complex, its deletion does not significantly impact transcriptional programs in either control B-cells, pre-tumoral Eµ-*myc* B-cells, or lymphomas: whether PRC1.6 is redundant altogether for transcriptional control or is involved in some other level of chromatin regulation in B-cells remains to be addressed.

Most importantly, our data do not formally rule out a role for Mga/Max and PRC1.6 in antagonizing Myc/Max-dependent transcription in DLBCL: it is indeed conceivable that, in the model used here, the Eµ-*myc* transgene might suffice to overcome the repressive function of Mga, while in DLBCL – or at least in a subset of cases – Mga may be critical to antagonize Myc activity and tumor progression, underlying the selective pressure to inactivate it (Reddy et al. 2017). Moreover, any of the five Mxd/Mnt proteins may also contribute repressive activity on common Mga- and Myc-target genes, and the balance between all these factors may differ between cell/tumor subtypes, experimental models and clinical cases. This notwithstanding, our data establish that, in conditions in which Mga shows no impact, Pcgf6 deletion clearly accelerates Myc-induced lymphomagenesis in the Eµ-*myc* mouse model.

The tumor suppressor activity of Pcgf6 reported here appears in contrast with the role of Pcgf4 (or Bmi1), a component of the canonical PRC1 complex (Scelfo et al. 2015) that has pro-tumoral activity in Eµ-*myc* mice, mediated by repression of the tumor suppressor locus *Cdkn2a* (or *Ink4/Arf*) (Jacobs et al. 1999). Most noteworthy here, Pcgf6 appears to antagonize the function of another canonical PRC1 subunit, Pcgf2, in anterior-posterior (A-P) specification during embryogenesis (Endoh et al. 2017). By analogy, the tumor suppressor activity or Pcgf6 might have been mediated by suppression of canonical PRC1 activity. However, our RNA-seq data did not support this hypothesis: *Cdkn2a* was expressed at very low levels in control B-cells and was de-repressed in pre-tumoral Eµ-myc B-cells, as expected (Eischen et al. 1999), but loss of Pcgf6 did not reverse this effect (Supplemental Fig. S3F). Together with its limited impact on global expression profiles, these observations suggest that Pcgf6 does not act by suppressing canonical PRC1 function during lymphomagenesis.

In conclusion, we have unraveled a distinct tumor suppressor activity of Pcgf6, unlinked from Mga and PRC1.6 - and possibly from any direct role in gene regulation. Besides the PRC1.6 complex, Pcgf6 interacts with the histone H3K4 demethylases JARID1c/d (Lee et al. 2007; Boukhaled et al. 2016) and may have alternative partners, yet to be investigated. Our findings warrant thorough characterization of alternative Pcgf6 activities and their relevance in human tumors.

## Materials and Methods

### Mouse strains and genotyping

Mice bearing the conditional allele *Mga*^*fl*^ (originally called *Mga*^*Inv*^) (Washkowitz et al. 2015) were bred with either CD19-*Cre* (Rickert et al. 1997) (a gift of Klaus Rajewsky) or *E*μ*-myc* transgenic animals (Adams et al. 1985), and the resulting compound mice bred to obtain the *Mga*-targeted cohort. The same strategy was pursued with the *Pcgf6*^*fl*^ allele (Endoh et al. 2017). The final crosses used to obtain our experimental cohorts are reported in Supplemental Table S1. Of note, the *Pcgf6*^*fl*^ cohort was inbred C57BL/6J, while the *Mga*^*fl/fl*^ cohort was of mixed genetic background (Washkowitz et al. 2015). In all experiments, gender and age-matched mice (both females and males) were used without randomization or blinding. Genomic DNA extraction and genotyping were performed as previously described (Bisso et al. 2020), with the PCR primers listed in Supplemental Table S5.

Experiments involving animals were done in accordance with the Italian Laws (D.lgs. 26/2014), which enforces Dir. 2010/63/EU (Directive 2010/63/EU of the European Parliament and of the Council of 22 September 2010 on the protection of animals used for scientific purposes) and authorized by the Italian Minister of Health with projects 391/2018-PR.

### Isolation and culturing of primary murine lymphoma cell lines

Mice were inspected personally for tumor development (Supplemental Materials and Methods). Infiltrated lymph nodes, spleen and bone marrow were collected and smashed in PBS. Cell suspensions were passed three times through a Falcon® 70 µm Cell Strainer (Corning, #352350), centrifuged (2000 rpm for 5 minutes) and resuspended in 10 ml of Erythrocyte Lysis buffer (150 mM NH_4_Cl, 10 mM KHCO_3_, 0,1 mM EDTA). After another centrifugation step, cells were resuspended in 10 ml of MACS buffer (PBS, 2mM EDTA, 0.5% BSA), and part of the cells used for *in vitro* culture. Primary cells were grown in suspension in B cell medium (BCM) composed of a 1:1 ratio of DMEM (Dulbecco’s Modified Eagle Medium, Euroclone, ECM0103L) and IMDM (Iscove’s Modified Dulbecco’s Medium, Sigma, I3390), supplemented with 10% fetal calf serum (Globefarm Ltd, Cranleigh, UK), 2 mM L-glutamine (Invitrogen Life Technologies, Paisley, UK), 1% non-essential amino acids (NEAA), 1% penicillin/streptomycin and 25 µM β-mercaptoethanol. A lymphoma cell line was considered as stabilized when the splitting ratio reached 1:10 every 2 days, which usually occurred upon 2 weeks of *in vitro* culture.

### Analysis of apoptosis, proliferation and surface markers

Apoptosis in bone-marrow derived B-cells was measured with the CaspGLOW™ Red Active Caspase Staining Kit (BioVision, #K190) following manufacturer’s guidelines. Proliferation was quantified by EdU staining: EdU (Invitrogen, #A10044) was dissolved in sterile PBC to a concentration of 5mg/ml; for *in vivo* proliferation studies, 1 mg EdU in a volume of 200 μl was injected intraperitoneally 2 hours before analysis, followed by staining with the 647 EdU Click Proliferation kit (BD Pharmingen™, #565456) according to manufacturer’s guidelines. Samples were stained with Hoechst DNA content dye, acquired on a FACSCelesta™ cytofluorimeter, and analyzed using FlowJo Version 10.4.0 software.

For staining of surface markers, cells were incubated in MACS buffer with fluorochrome-conjugated antibodies (used at the dilutions indicated in Supplemental Table S5) for at least 1 hour at 4°C in the dark, and analysed by flow cytometry, as above.

### Statistical analysis

All experiments were performed at least in biological triplicates. Sample size was not predetermined but is reported in the respective Figure legends. P-values were calculated with One-way Anova using Tukey correction, except in the Fig.1A for Kaplan-Meier survival curves where log p-rank test was used.

## Data availability

The RNA-seq data produced in this work have been deposited in NCBI’s Gene Expression Omnibus (https://www.ncbi.nlm.nih.gov/geo/) and are accessible though the GEO Series accession number GSE190000.

## Competing interest statement

The authors declare no competing interest.

## Acknowledgments

We thank Stefano Campaner, Francesco Nicassio, Diego Pasini and members of the Amati lab for discussions, insight, suggestions and reagents, A. Gobbi, M. Capillo and all members of the animal facility for their help with the management of mouse colonies, S. Bianchi, L. Rotta, T. Capra and L. Massimiliano for assistance with Illumina sequencing, S. Ronzoni for assistance with flow cytometry, M.G. Jodice, F. Montani, G. Bertalot and S. Pece for the help with processing tissue samples, V.E. Papaioannou for providing *Mga*^*fl/fl*^ mice, and K. Rajewsky for CD19-*Cre* mice. This work was supported by grants from the Italian Health Ministry (RF-2011-02346976), from the Italian Association for Cancer Research (AIRC, IG2015-16768 and IG2018-21594) to B. Amati, and from the Ministry of Education, Culture, Sports, Science and Technology of Japan (Grants-in-Aid for Scientific Research, 23249015) to H. Kozeki.

## Authors’ Contributions

NT and AB designed and performed most of the experiments. M. Dalsass and MF performed bioinformatic data analysis. GC contributed to the *in vivo* experiments. AV, M. Doni and PN provided technical support, and DO the pathological analyses. HK and DP provided the *Pcgf6* mutant mice. AS helped with analysis and interpretation of RNA-seq and ChIP-seq experiments. NT, AB and BA wrote the manuscript. BA and AB conceived the project and co-supervised the work.

## Notes

https://www.ncbi.nlm.nih.gov/geo/query/acc.cgi?acc=GSE190000

### Competing Interest Statement

The authors have declared no competing interest.

