## Supplemental Information for "Polycomb Group Ring Finger Protein 6 suppresses Myc-induced lymphomagenesis"

for

**Contents:** **Page:**

**Supplemental Materials and Methods** 2

**Supplemental References** 4

**Supplemental Figures** 5

**Supplemental Tables** 10

**Supplemental Materials and Methods**

*Mouse monitoring and handling*

Eμ-*myc* transgenic mice were monitored 2-3 times a week for tumor development by visual inspection and peripheral lymph node palpation, and were sacrificed as soon as they showed signs of lymphoma (i.e. enlarged lymph nodes) (Adams et al. 1985). For pre-tumoral analysis, mice were collected at 4-6 weeks of age: spleen and bone marrow were dissected and processed for molecular analysis as previously described (Campaner et al. 2010).

*Immunoblotting*

Protein extraction and immunoblotting were performed as previously described (Bisso et al. 2020) with the indicated primary antibodies (Supplemental Table S5).

*Hematoxylin and Eosin staining*

For Hematoxylin and Eosin staining and pathological analysis tissues were collected and processed as follows. Freshly isolated lymphoma samples were washed in PBS, fixed in 4% (v/v) paraformaldehyde at 4°C degrees for at least 16-24 hours, washed in PBS, and stored in 70% ethanol at 4°C for a maximum of 1 week before inclusion. For the latter, each tissue was dehydrated with increasing concentrations of ethanol, embedded in paraffin blocks and stored at RT. For Hematoxylin and Eosin staining each block was cut into 3/5-mm thick sections and mounted on glass slides. Slides were counterstained with Harris Hematoxylin (Sigma-Aldrich, #HHS80) and Eosin Y solution (Sigma-Aldrich, #HT110216), dehydrated through alcoholic scale, and mounted with Eukitt (Bio-Optica, #09-00250). All images were acquired with the Aperio Digital Pathology Slide Scanner ScanScopeXT (Leica) prior to pathological evaluation.

*RNA sequencing*

RNA extraction, processing and sequencing, as well as the filtering of RNA-seq reads and bioinformatic and statistical analyses, were performed as previously described (Tesi et al. 2019; Bisso et al. 2020; Pellanda et al. 2021). The analysis of tumor clonality from RNA-seq reads was performed as previously described (Barbosa et al. 2020). Sequences of the PCR primers used are reported in Supplemental Table S5.

*ChIP sequencing*

The fixation of *in vitro* stabilized lymphoma cell lines and their processing for chromatin immunoprecipitation (ChIP) was performed as previously described (Sabò et al. 2014). 5 µg of each of the antibodies listed in in Supplemental Table S5 were used to immuno-precipitate either 500 µg (for the mapping of Myc, Max and Pcgf6) or 250 µg of fixed chromatin (for the histone marks H3K4me3, H3K4me1, H3K27ac, H3K27me3 and H2Ak119Ub). While Myc and Max precipitates were processed exactly as in Sabò et al. (2014), Pcgf6 and histone mark precipitates were processed as in Scelfo et al. (2019). 1.5-2 ng of DNA was then used to generate the chromatin immunoprecipitation sequencing (ChIP-Seq) libraries according to the Illumina protocol, and sequenced with the Illumina NovaSeq 6000.

ChIP-seq reads were analyzed as previously published (Sabò et al. 2014; Pellanda et al. 2021). Peaks were mapped and annotated according to the genomic position of their midpoint, as (i.) promoter: between -2Kb and +1Kb from the annotated refgene start coordinate or transcriptional start site (TSS); (ii.) gene body: between > 1Kb from the TSS to the 3' end of an annotated refgene; (iii.) intergenic: all peaks positioned outside of the aforementioned intervals. Qualitative and quantitative heatmaps of ChIP-seq enrichment were generated using R with Bioconductor and compEpiTools packages, tools for computational epigenomics (Gentleman et al. 2004; Kishore et al. 2015).

*Oligonucleotide Primers*

Primers for mRNA analysis were designed with Primer-BLAST (https://www.ncbi.nlm.nih.gov/tools/primer-blast/) (Ye et al. 2012). The complete list of primers used in this study is shown in Supplemental Table S5.

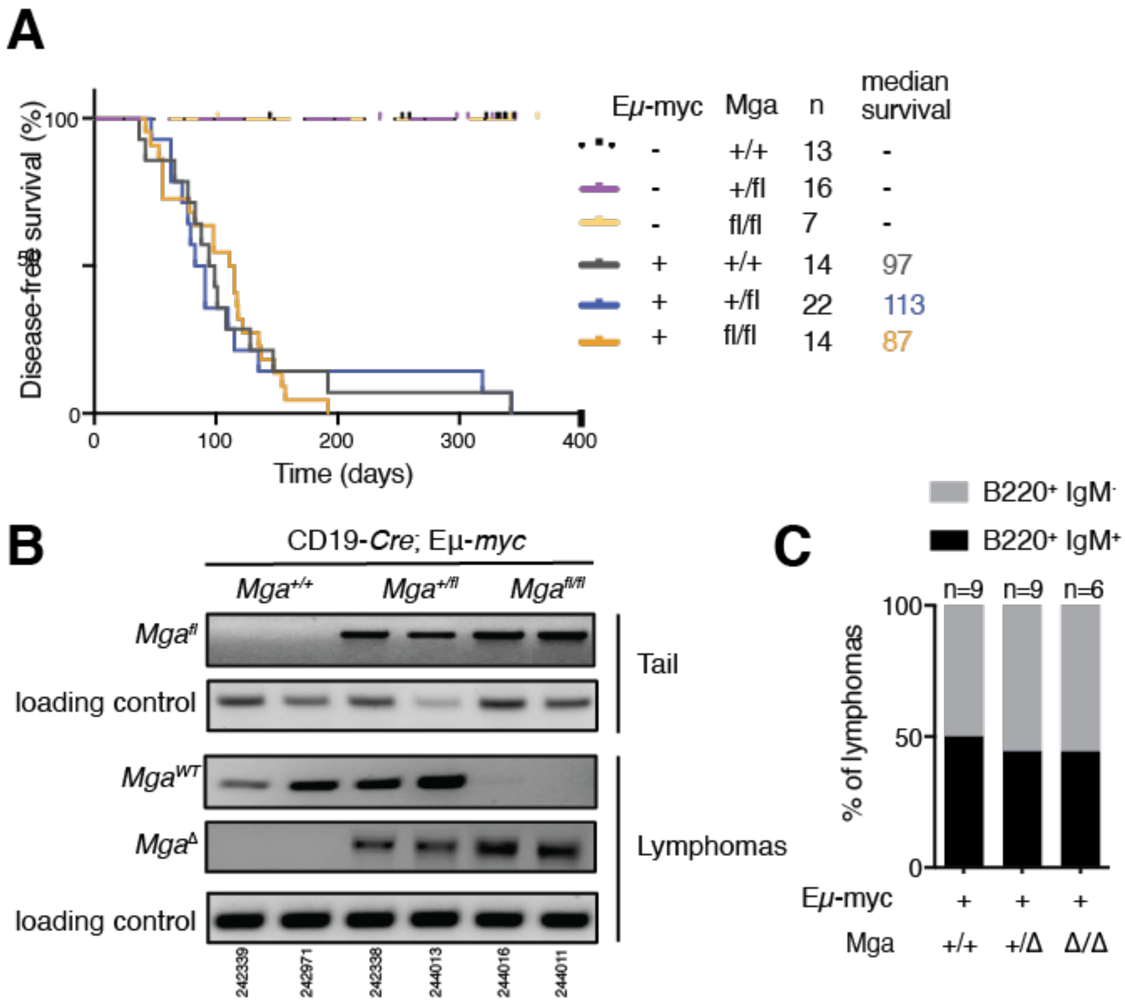


**Supplemental Figure S1. Characterization of *Mga*-mutant lymphomas. A.** Disease-free survival curves for mice of the indicated Eµ-*myc* and *Mga* genotypes (all with the CD19-*Cre* transgene). The number of mice (n) and the median survival (in days) are indicated. Of note, control Eµ-*myc;Mga*$^{+/+}$ mice showed faster lymphoma onset relative to their Eµ-*myc;Pcgf6*$^{+/+}$ counterparts (Fig. 1A), owing most likely to the different genetic backgrounds of the two cohorts (mixed and inbred C57BL/6J, respectively). **B.** Recombination status of the *Mga*$^{fl}$ allele in lymphomas, as determined by semi-quantitative PCR on 10 ng of genomic DNA isolated from sorted CD19+ tumor cells. Tails from the same mice were used as a negative control. Two mice per genotype are represented, with their IDs indicated at the bottom. Allele-specific PCR reactions were performed with the primers listed in Supplemental Table S5. *Mga*$^{fl}$: non recombined allele; *Mga*$^{\Delta}$: recombined allele; *Mga*$^{WT}$: wild type allele. **C.** Immunophenotyping of B220 and IgM revealed similar proportions of B220$^{+}$ IgM$^{+}$ and B220$^{+}$ IgM$^{-}$ tumors among Eµ-*myc* lymphomas of the indicated *Mga* genotypes. The numbers above each bar represent number of mice analyzed for each genotype.

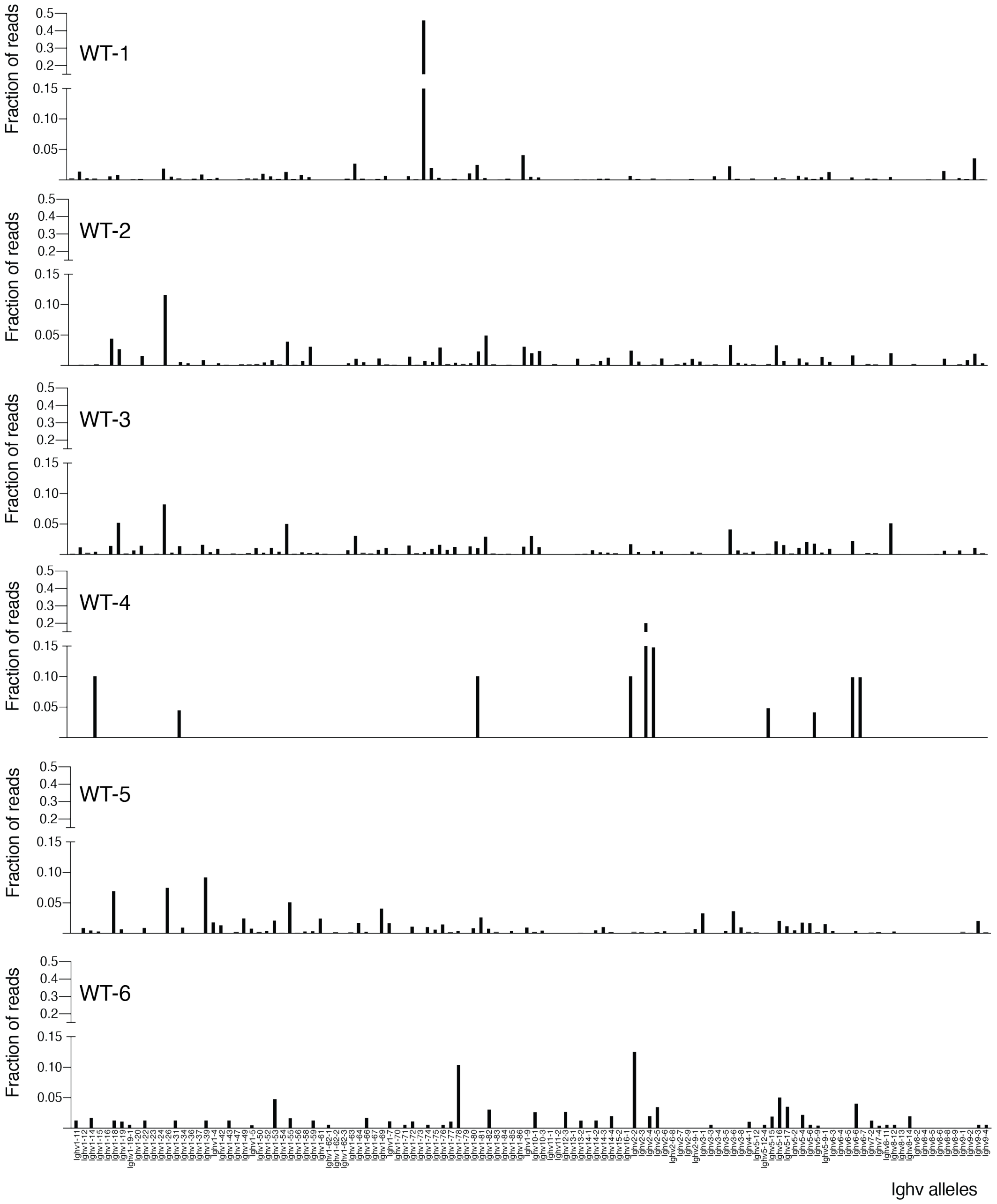


**Supplemental Figure S2.** Continued on the next page.

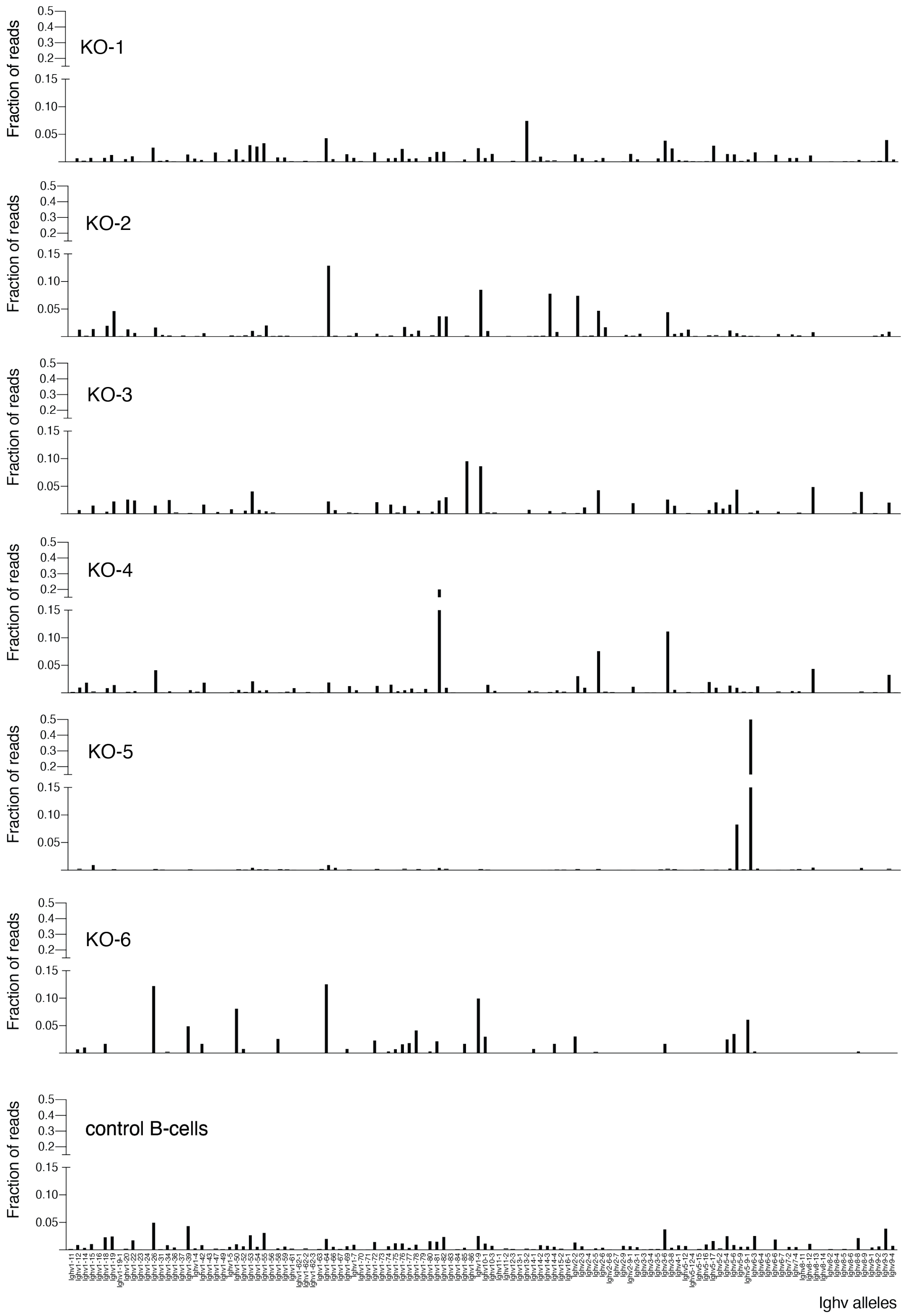


**Supplemental Figure S2. Characterization of clonality of *Pcgf6*-mutant lymphomas.** The graphs show the fraction of RNA-seq reads mapped to each individual IgH-V allele (displayed in the same order as listed in Supplemental Table S3) relative to the total reads mapped to all IgH-V alleles. Histograms for each sample analyzed – Eμ-*myc; Pcgf6*$^{+/+}$ lymphomas (n=6); Eμ-*myc*; *Pcgf6*$^{\Delta/\Delta}$ lymphomas (n=5); control B-cell

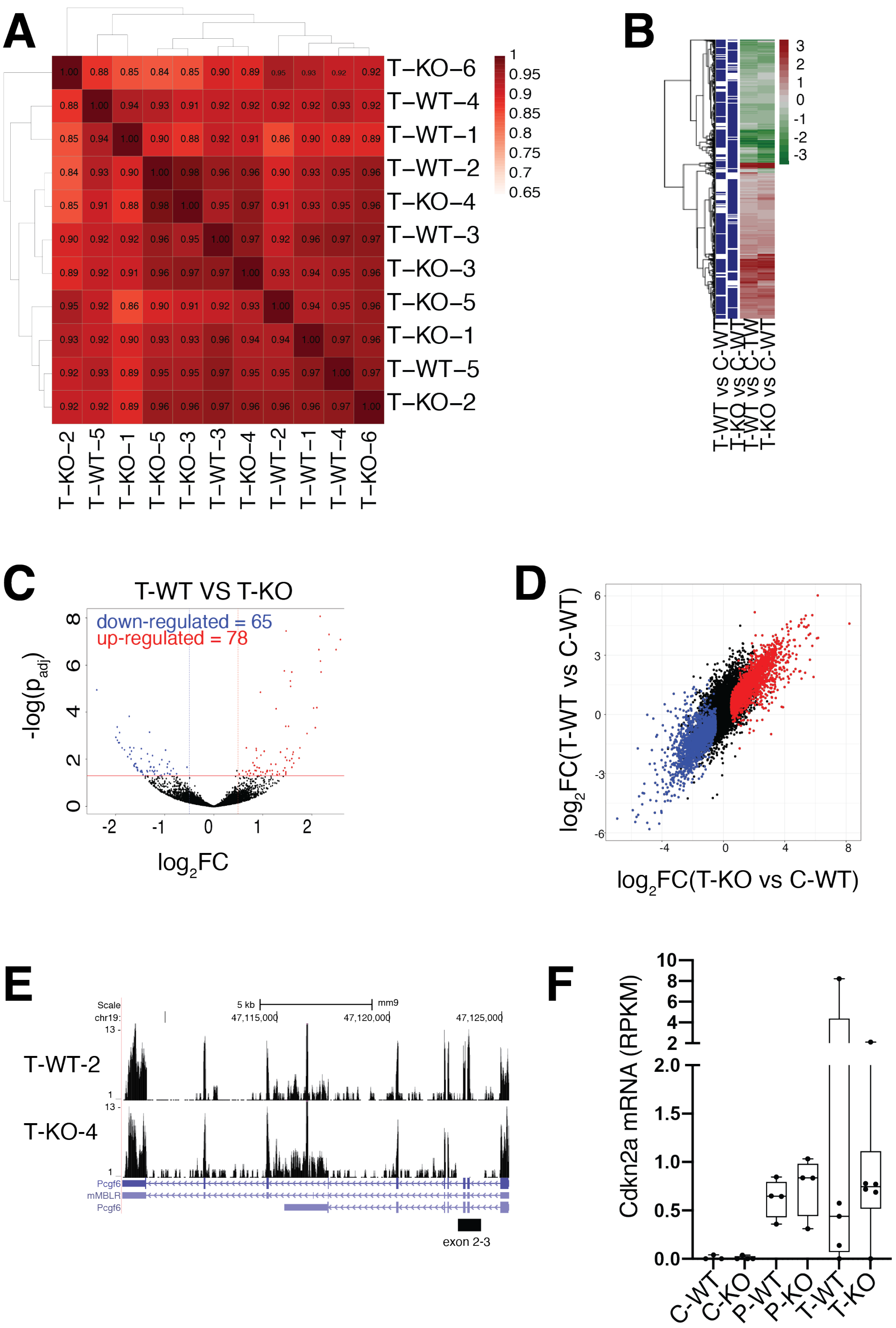


**Supplemental Figure S3. Pcgf6 loss does not affect Myc-dependent transcription.** RNA-seq profiles were generated from tumor (T) samples of either *Pcgf6* genotype (WT and KO): samples are labeled as defined in Fig. 3A. **A.** Pearson correlation between tumor samples, based on their RNA-seq profiles. **B.** Clustered heatmap of differentially expressed genes (DEGs) in tumoral samples. The two left columns show a blue line in correspondence of a gene identifies as a DEG (q <0.05); the two right columns show the $log_2FC$ value for each DEG colored from green (-3) to red (3). **C.** Same as Fig. 3A, showing the DEGs called between the T-KO and T-WT samples. **D.** Comparison of the DEGs called in T-WT (Y-axis) and T-KO (X-axis), both relative to C-WT. The DEGs are colored based on the call in the x-axis. **E.** Representative RNA-seq profiles along the *Pcgf6* locus shown in the UCSC Genome Browser, from the indicated WT and KO tumors. Note the absence of reads in the targeted exon 2-3 region (indicated below the map) in the KO sample. **F.** *Cdkn2a* mRNA levels, as determined by RNA-seq, normalized to the C-WT samples.

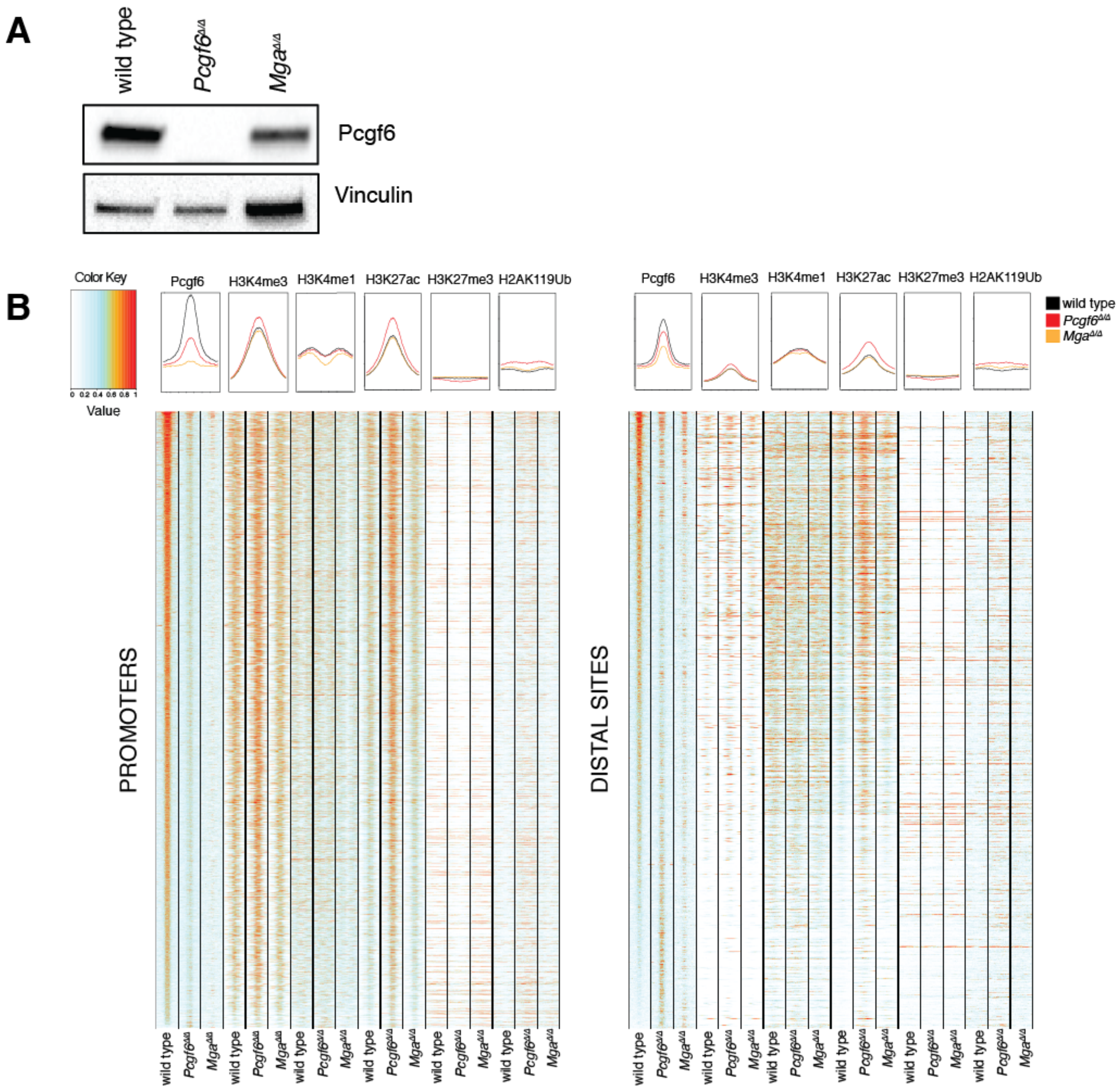


**Supplemental Figure S4. Pcgf6 is binding preferentially active promoters in B-cell lymphomas. A.** Immunoblot analysis of Pcgf6 in wild type, *Pcgf6*$^{\Delta/\Delta}$ and *Mga*$^{\Delta/\Delta}$ lymphomas. Vinculin was used as loading control. Note that we were unable to detect the Mga protein in mouse samples so far, owing to the lack of adequate antibodies. **B.** Heatmaps representing normalized ChIP-seq intensity for Pcgf6 (ranked by decreasing intensity, from top to bottom) and the indicated histone marks in the same lymphomas. Left: the heatmap includes every annotated promoter that was called as Pcgf6-associated by ChIP-seq in at least one of the samples; each column spans the interval between -2 and +2 kb from the TSS. Right: includes all Pcgf6-associated distal sites, spanning the interval between -2 and +2 kb from the center of the Pcgf6 peak.

**A. Breeding:** ***CD19-Cre/Cre; Pcgf6^+/fl^* x *Eμ-myc; Pcgf6^+/fl^***

| Genotype | Expected frequency | Observed frequency | |
|---|---|---|---|
| CD19-*Cre; Pcgf6*$^{+/+}$ | 12.50% | 21/173 | 12.14% |
| CD19-*Cre; Pcgf6*$^{+/fl}$ | 25% | 52/173 | 30.06% |
| CD19-*Cre; Pcgf6*$^{fl/fl}$ | 12.50% | 30/173 | 17.34% |
| CD19-*Cre*; Eμ-*myc*; *Pcgf6*$^{+/+}$ | 12.50% | 36/173 | 20.81% |
| CD19-*Cre*; Eμ-*myc*; *Pcgf6*$^{+/fl}$ | 25% | 29/173 | 16.76% |
| CD19-*Cre*; Eμ-*myc*; *Pcgf6*$^{fl/fl}$ | 12.50% | 5/173 | 2.89% |

**B. Breeding:** ***CD19-Cre; Mga*$^{+/fl}$ x *Eμ-myc; Mga*$^{+/fl}$**

| Genotype | Expected frequency | Observed frequency | |
|---|---|---|---|
| *Mga*$^{+/+}$ | 6.25% | 6/96 | 6.25% |
| *Mga*$^{+/fl}$ | 12.50% | 23/96 | 23.96% |
| *Mga*$^{fl/fl}$ | 6.25% | 2/96 | 2.08% |
| Eμ-*myc*; *Mga*$^{+/+}$ | 6.25% | 7/96 | 7.29% |
| Eμ-*myc*; *Mga*$^{+/fl}$ | 12.50% | 3/96 | 3.13% |
| Eμ-*myc*; *Mga*$^{fl/fl}$ | 6.25% | 9/96 | 9.38% |
| CD19-*Cre*; *Mga*$^{+/+}$ | 6.25% | 11/96 | 11.46% |
| CD19-*Cre*; *Mga*$^{+/fl}$ | 12.50% | 19/96 | 19.79% |
| CD19-*Cre*; *Mga*$^{fl/fl}$ | 6.25% | 3/96 | 3.13% |
| CD19-Cre; Eμ-*myc*; *Mga*$^{+/+}$ | 6.25% | 4/96 | 4.17% |
| CD19-Cre; Eμ-*myc*; *Mga*$^{+/fl}$ | 12.50% | 5/96 | 5.21% |
| CD19-Cre; Eμ-*myc*; *Mga*$^{fl/fl}$ | 6.25% | 4/96 | 4.17% |

**Supplemental Table S1. Breeding Strategy.** Each table shows the expected (assuming mendelian distribution) and observed frequencies of the indicated compound genotypes, based on the crosses shown at the top. **A**. *Pcgf6* mutant cohort. Here all siblings are positive for CD19-*Cre*, as this transgene was first bred to homozygosity in one of the parents (CD19-*Cre/Cre*). Note that Eμ-*myc* and *Pcgf6* segregated in a sub-Mendelian manner ($p < 0.0001$), consistent with their close genomic location on chromosome 19 (Lefebure et al. 2017) (http://www.informatics.jax.org/marker/MGI:1918291). **B**. *Mga* mutant cohort. In line with published data (Washkowitz et al. 2015), *Mga*$^{fl/fl}$ mice were recovered at sub-Mendelian frequencies (p-value $< 0.005$), confirming that *Mga*$^{fl}$ is a hypomorphic allele.

**Supplemental Tables S2-S4.** These tables are provided as separate Excel files and include the following:

- **Supplemental Table S2. Pathological Analysis of Eμ-myc *Pcgf6*-mutant lymphomas.** Detailed pathological analysis and resulting tumor classification as defined for the human disease (Swerdlow et al. 2016) for the indicated mouse lymphomas, identified by their Genotype and Sample ID.
- **Supplemental Table S3. Analysis of tumor clonality.** The B-cell clonality of the indicated samples was determined based on the fraction of reads for each Ighv allele in RNA-seq data, as previously described (Barbosa et al. 2020). To determine this value, we first scored the RPM (reads per million) for each allele: read counts were first normalized based on exon length (to account for different exon lengths), then divided by the total number of Igvh reads (to compensate for a different read depth in each sample), and finally multiplied by 1 million. The fraction of reads was determined by dividing the RPM value by the sum of the RPMs of all Igvh alleles.
- **Supplemental Table S4. Differentially Expressed Genes (DEGs) called in this work.** The first spreadsheet (All DESeq2) shows DESeq2 calculations of log2FC and padj for all genes, in each or the indicated pairwise comparisons (Sample pairs). The additional spreadsheets show the DEGs called in each comparison (padj<0.05).

**A. Antibodies for immunoblotting**

| **Antibody** | **Source** | **Dilution/mg used** |
|---|---|---|
| Myc Y69 | Abcam (ab32072) | 1:2000 |
| Vinculin | Sigma-aldrich (V9264) | 1:10000 |
| Max | Bethyl (A302-866A) | 1:1000 |
| Pcgf6 | Scelfo et al. 2019 | 1:1000 |
| Hsp90 | Santa Cruz (sc-13119) | 1:10000 |

**B. Antibodies for flow cytometric analysis**

| **Antibody** | **Conjugate** | **Company** | **Clone** | **Dilution used** |
|---|---|---|---|---|
| B220 | PE | BD Pharmingen (#553089) | RA3-6B2 | 1:200 |
| B220 | efluor 450 | eBioScience (48-0452-82) | RA3-6B2 | 1:200 |
| CD19 | PE-Cy7 | BD Pharmigen (552854) | 1D3 | 1:400 |
| CD21 | PE | eBioScience (12-0211-82) | 8D9 | 1:800 |
| CD23 | AlexaFluor 647 | BioLegend (101612) | B3B4 | 1:100 |
| CD25 | APC | eBioScience (17-0251-82) | PC61.5 | 1:100 |
| CD43 | FITC | eBioScience (11-0431-85) | R2/60 | 1:200 |
| IgD | BV510 | BD Pharmigen (#563110) | 11-26C.2A | 1:200 |
| IgM | APC | eBioScience (47-5790-82) | II/41 | 1:200 |
| IgM | APC-Cy7 | BioLegend (406516) | RMM-1 | 1:200 |
| CD45 | BV786 | BD Pharmingen (#564225) | 30-F11 | 1:!00 |

**Supplemental Table S5.** List of materials used in this work *(continued on the next page).*

### C. Antibodies for chromatin immunoprecipitation.

| Antibody | Company |
|---|---|
| Max | Bethyl (A302-866A) |
| Pcgf6 | Scelfo et al. 2019 |
| Myc N262 | Santa Cruz (sc-764) |
| H3K4me3 | Active Motif (#39159) |
| H3K4me1 | Abcam (ab8895) |
| H3K27ac | Abcam (ab4729) |
| H3K27me3 | Cell Signaling (#9733) |
| IgG | Santa Cruz (sc-2027) |
| H2AK119Ub | Cell Signaling |

### D. PCR Primers. D. PCR Primers.

| Application | Locus | Forward primer | Reverse primer | Amplicon (bp) |
|---|---|---|---|---|
| **Expression (RT-PCR)** | mouse Pcgf6 | CTTCTCTCTGCGTCTGGAGTC | TCAGCTCGACAAGGTTTATCAG | 77 |
| | mouse c-Myc | TTTTTGTCTATTTGGGGACAGTG | CATCGTCGTGGCTGTCTG | 130 |
| | mouse Max | CCTGGGCCGTAGGAAATGAG | CAGCCGCAGATTGAAACCTC | 82 |
| | mouse Mga | AAATCTTTAACTGCTGCCAAGAA | CTGCAACCTGAATCATTTGTGGT | 200 |
| | mouse H3 | GTGAAGAAACCTCATCGTTACAGGCCTGGT | CTGCAAAGCACCAATAGCTGCACTCTGGAA | 177 |
| **Analysis of recombination on gDNA by semi-quantitative PCR** | Mga$^{wt}$ | ATTCCTGTAGGCCCTGGAAG | GGGAGGATTGGGAAGACAAT | 325 |
| | Mga$^{null}$ | ATTCCTGTAGGCCCTGGAAG | CAGGACAACCTGACACCTCTG | 600 |
| | Internal Positive Control (IL-2) | CTAGGCCACAGAATTGAAAGATCT | GTAGGTGGAAATTCTAGCATCATCC | 324 |
| **Genotyping** | CD19 wt | CCAGACTAGATACAGACCAG | AACCAGTCAACACCCTTCC | 452 |
| | CD19-*Cre* | CCAGACTAGATACAGACCAG | TCAGCTACACCAGAGACGG | 750 |
| | Eμ-*myc* | GGTTTAATGAATTTGAAGTTGCCA | TTCTTGCCCTGCGTATATCAGTC | 210 |
| | Mga$^{wt}$ | CAGGACAACCTGACACCTCTG | GGTATGGTTGTAATGATCAGCTTTC | 325 |
| | Mga$^{Inv}$ | CAGGACAACCTGACACCTCTG | GCTGGGGCTCGATCCTCTAG | 500 |
| | Pcgf6$^{wt}$ | TTAATTGCTGCGTTCCATCTC | ATGTCAGAGAACTGGGACCGC | 404 |
| | Pcgf6$^{fl}$ | TTAATTGCTGCGTTCCATCTC | GGCTAGATCTGCTGGAGACTT | 378 |

**Supplemental Table S5.** List of materials used in this work.
